# The atypical Rho GTPase RhoU interacts with Intersectin-2 to regulate endosomal recycling pathways

**DOI:** 10.1101/631440

**Authors:** Olga Gubar, Pauline Croisé, Sergii Kropyvko, Tetyana Gryaznova, Petra Toth, Anne Blangy, Nicolas Vitale, Alla Rynditch, Stéphane Gasman, Stéphane Ory

## Abstract

Rho GTPases play a key role in various membrane trafficking processes. RhoU is an atypical small Rho GTPase related to Rac/Cdc42 which possesses unique N- and C-terminal domains that regulate its function and its subcellular localization. RhoU localized at the plasma membrane, on endosomes and in cell adhesion structures where it governs cell signalling, differentiation and migration. However, despite its endomembrane localization, RhoU function in vesicular trafficking has been unexplored. Here, we identified intersectins (ITSNs) as new binding partners for RhoU and showed that the second PxxP motif at the N-terminus of RhoU mediated interactions with SH3 domains of ITSNs. To evaluate the function of RhoU and ITSNs in vesicular trafficking, we used fluorescent transferrin as a cargo for uptake experiments. We showed that silencing of either RhoU or ITSN2, but not ITSN1 increased transferrin accumulation in early endosomes resulting from defect in fast vesicle recycling. Concomitantly, RhoU and ITSN2 colocalized to a subset of Rab4-positive vesicles suggesting that RhoU-ITSN2 interaction may occur on fast recycling endosomes to regulate the fate of vesicular cargos.

## Introduction

Rho GTPases are small G proteins belonging to the Ras superfamily. They regulate many cell functions including cell migration, cell polarity and cell proliferation (Jaffe and Hall, 2005). Although most studies have been focused on the role of the canonical members of the family namely RhoA, Rac1 and Cdc42, evidences emerged for critical functions of the so-called atypical Rho GTPases which encompass Rnd, RhoH, RhoU, RhoV and RhoBTB proteins (Aspenström et al., 2007). Atypical Rho GTPases are mostly found associated to membranes and in a constitutive active state due to high guanine nucleotide exchange rate (RhoU, RhoV) or due to amino acids substitutions that render them defective for GTPase activity (Rnd proteins, RhoH). Activities of atypical Rho GTPases are primarily regulated by the balance between transcription/translation, degradation and binding to regulatory proteins. Indeed, compared to canonical members, most atypical Rho GTPases possess additional domains at their N- and/or C-terminus that bind various proteins and contribute to regulate their activities (Aspenström et al., 2007; Vega and Ridley, 2008).

The atypical Rho GTPase RhoU was first isolated as a gene transcriptionally up-regulated in wnt-1 transformed mouse mammary epithelial cells (Tao et al., 2001). RhoU shares significant homology with Cdc42 as well as some biological functions. For example, RhoU binds and activates the p21 activated kinase (PAK1), induces filopodia and regulates cell tight junctions (Brady et al., 2009; Garrard et al., 2003; Saras et al., 2004; Tao et al., 2001). In addition, amino acids of the effector loop that mediate binding to downstream targets such as PAK1 or Par6 are conserved between RhoU and Cdc42 (Brady et al., 2009; Ory et al., 2007). However, RhoU possesses N- and C-terminal extensions which confer unique properties. The N-terminal extension enriched in proline residues mediates binding to SH3 domains possessing proteins such as Nck1 and Grb2 (Saras et al., 2004; Shutes et al., 2004). Its deletion increases RhoU transforming activity as well as RhoU signaling through PAK1 activation (Berzat et al., 2005; Brady et al., 2009). Mutations of proline-rich motifs of RhoU instead impaired EGF receptor signaling by preventing RhoU recruitment by Grb2 to EGF receptor complex (Zhang et al., 2011). Whereas its activity appeared to be regulated by its N-terminal extensions, RhoU subcellular localization is mostly controlled by its 21 amino acids long C-terminal extension which ends up with a non conventional CAAX box (Berzat et al., 2005) and its effector loop (Ory et al., 2007). RhoU has been localized at the plasma membrane, on endosomes and at focal adhesions where it regulates their turnover (Chuang et al., 2007; Ory et al., 2007). RhoU association with membranes is mostly dependent on its CAAX motif (Berzat et al., 2005; Zhang et al., 2011) whereas a bipartite motif comprising the C-terminal extension and the effector loop is needed for focal adhesion targeting (Ory et al., 2007). Collectively, these studies revealed that the N- and C-terminal extension of RhoU have distinct function, the first one regulating its activity and the second one contributing to its subcellular localization.

To further characterize RhoU and its function, we searched for new binding partners by pull-down experiments and found that Intersectins (ITSNs) formed a complex with RhoU. As ITSNs are multidomain endocytic proteins mostly involved in clathrin-mediated endocytosis pathways (Tsyba et al., 2011), we next investigated the importance of RhoU and ITSNs in transferrin trafficking pathway and found that RhoU and ITSN2 coordinate the recycling of transferrin receptors.

## Results

### ITSN1 and ITSN2 interact with RhoU

Preliminary data based on GST-RhoU pull down experiments combined to mass spectrometry analysis suggested that the short form of ITSN2 (ITSN2-S), a member of the intersectin (ITSN) family could interact with RhoU (Ory S. and Blangy A., unpublished data). Moreover, bioinformatics analysis for potential RhoU binding motif using Scansite database (http://scansite3.mit.edu; Obenauer et al., 2003), picked the SH3A domain of ITSN1 with the highest score whereas the SH3 domain of Nck, a known partner of RhoU (Shutes et al., 2004) came second (supplementary Figure S1). This potential interaction between RhoU and ITSN isoforms was further investigated using GST pull down and immunoprecipitation experiments (Fig.1). Intersectin proteins are encoded by two genes, *itsn1* and *itsn2*, and both proteins exist in at least one short (S) and one long (L) forms resulting of alternative splicing (Fig.1A)

**Figure 1:**
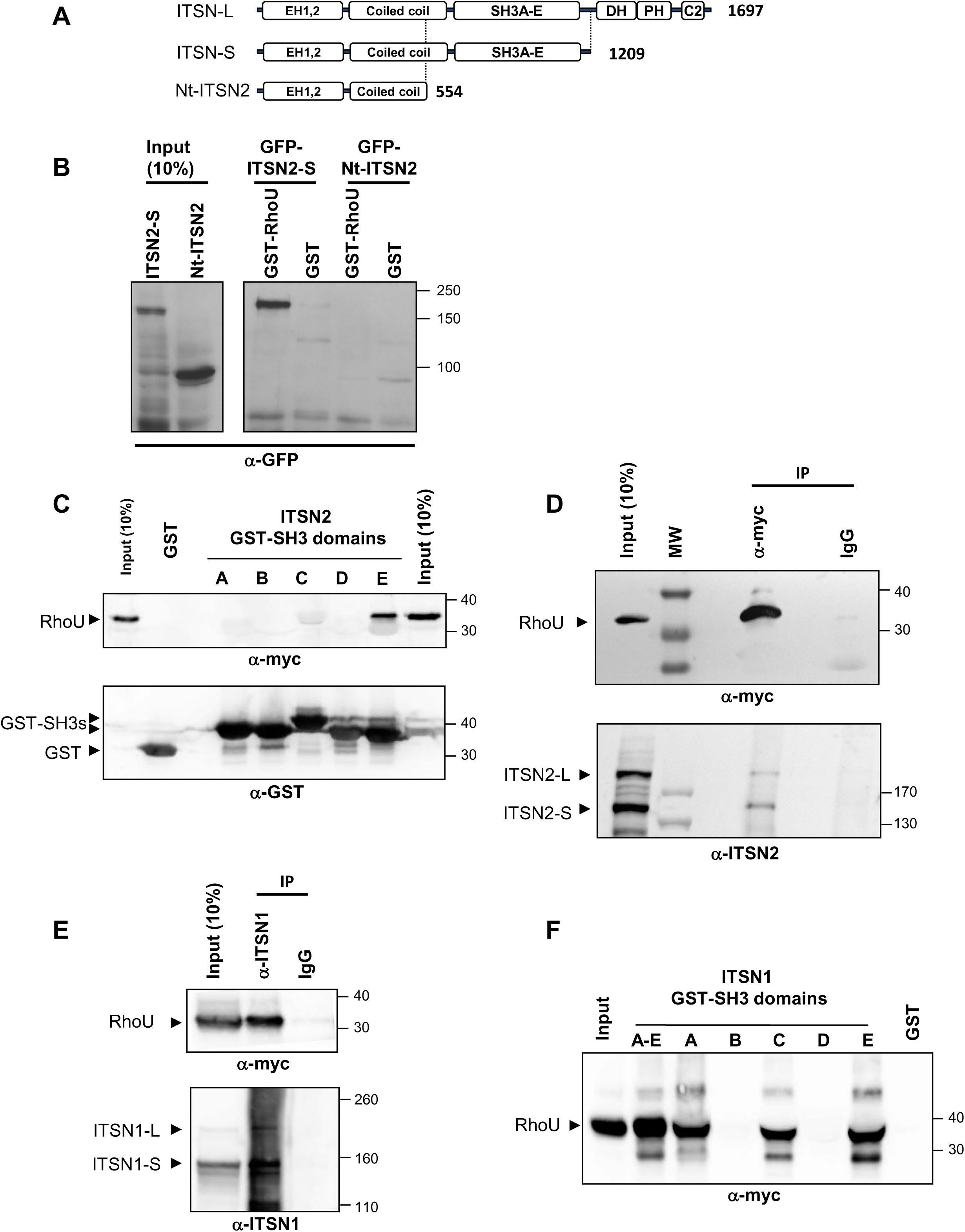
RhoU interacts with ITSN proteins. **(A)** Schematic representation of the intersectin isoforms. Both short (ITSN-S) and long (ITSN-L) forms contains two Epsin homology domains (EH1, EH2), a coiled-coil domain and five Src homology 3 domains (SH3A-E) whereas only the long form (ITSN-L) contains three additional domains, a Dbl homology domain (DH) catalyzing the activation of Cdc42, a Pleckstrin homology domain (PH) needed for plasma membrane targeting and a C2 domain. Nt-ITSN2 contains EH domains and a truncated coiled coil domain. Number indicates amino acid length of encoded proteins. **(B)** HEK 293T cells were transfected with GFP-ITSN2-S or Nt-ITSN2 and lysates subjected to pull down experiments using either recombinant GST or GST-WT RhoU. Precipitated proteins were revealed by western blot using anti-GFP antibodies. **(C)** Pull down experiments using individual SH3 domain (SH3A to SH3E) of ITSN2 fused to GST and lysates of HEK 293T expressing myc-RhoU. Precipitated RhoU was revealed by western blot with anti-myc antibodies and GST fusion proteins with anti-GST antibodies. **(D)** Immunoprecipitation (IP) experiments from lysates of cells expressing myc-tagged RhoU using anti-myc antibodies. Arrowheads point to both form of precipitated endogenous ITSN2 (S+L). **(E)** Immunoprecipitation of endogenous ITSN1 (S+L) from HEK293T cells transfected with myc-tagged RhoU. Precipitated RhoU was revealed by anti-myc antibodies. Arrowheads point to both forms of ITSN1. **(F)** HEK293T cells were transfected with myc-tagged WT RhoU and cell lysates incubated with GST alone, GST fused to individual SH3 domain (SH3A to SH3E) or to the five SH3 domains (SH3A-E) of ITSN1. Precipitated RhoU was revealed by western blot with anti-myc antibodies.

To validate the potential formation of a complex between RhoU and ITSN2-S, we first performed pull down experiments using purified recombinant GST-RhoU with cell extracts from HEK293T cell lines expressing GFP-ITSN2-S. Western blot analysis using anti-GFP antibodies revealed that ITSN2-S precipitated with GST-RhoU (Fig.1B). As a truncated mutant of ITSN2-S deleted of its SH3 domains (Nt-ITSN2, Fig.1A) was not precipitated by GST-RhoU, the formation of a complex between RhoU and ITSN2 is most likely mediated through one or several SH3 domains (Fig.1B). To determine which one of the five SH3 domains of ITSN2 could be involved, pull down experiments were performed using individual SH3 domain fused to GST. As illustrated in Figure 1C, only the GST-SH3-E domain precipitated myc-RhoU from HEK293T cell extracts suggesting that RhoU binds specifically to the SH3E domain of ITSN2 and not to the others. To further demonstrate that RhoU-ITSN2 interaction occurs in cells, we performed immunoprecipitation experiments in HEK293T cells expressing myc-RhoU. Both endogenous ITSN2-S and 2-L were co-immunoprecipitated by myc-RhoU suggesting that both forms of ITSN2 were associated with RhoU in cells (Fig.1D). Moreover, in line with our database analysis suggesting that RhoU possesses a potential ITSN1 binding site, ITSN1-S and ITSN1-L also co-precipitated with myc-RhoU in HEK293T cells demonstrating that RhoU can interact with both ITSN1 and ITSN2 (Fig.1E).

Finally, GST pull down experiments using the five (SH3A-E) or individual SH3 domains of ITSN1 showed that SH3-A, -C, and –E efficiently precipitated RhoU whereas SH3-B and -D did not (Fig.1F), suggesting that the SH3 domains also mediated the interaction between ITSN1 and RhoU. Altogether, these in vitro experiments indicated that specific SH3 domains of ITSN1 and ITSN2 form a complex with RhoU.

### N-terminal proline-rich domains of RhoU mediate binding to SH3 domains of ITSNs

RhoU has been classified as an atypical Rho protein notably because of additional domains found at its N- and C- terminal regions compared to its close homolog Cdc42 (Aspenström et al., 2007; Shutes et al., 2004). RhoU N-terminal extension possesses two PxxPxR sequences corresponding to consensus SH3 binding motifs (Shutes et al., 2004; Yu et al., 1994). To identify which proline-rich motif of RhoU is important for binding to ITSN SH3 domains, we expressed RhoU mutants in which proline to alanine substitutions were introduced into individual motif (M1 or M2) or into both motifs (DM, Fig.2A). These mutations have been previously shown to disrupt RhoU binding to SH3 domains of the adaptor protein Grb2 (Zhang et al., 2011). We performed pull down assays with SH3A-E ITSN1 on cell lysates of HEK293T cells expressing myc-RhoU and found that M1 mutations only slightly reduced RhoU binding whereas M2 and DM mutations completely abolished it indicating that the second proline-rich motif of RhoU is the main binding site for ITSN1 SH3 domains (Fig.2B). To confirm that the proline-dependent interaction occurred also in cells, the myc-RhoU mutants were co-expressed with Omni-tagged ITSN1 in HEK293T for co-immunoprecipitation experiment. In agreement with pull down experiments, amounts of ITSN1 co-immunoprecipitated with RhoU were dramatically reduced with the M2 and DM mutants whereas M1 mutations had only a modest effect (Fig.2C). Similar pull down experiments were performed using the SH3A-E domains of ITSN2 showing that M2 and DM mutants were unable to bind ITSN2 SH3 domains (Fig.2D). Altogether, these data indicate that the second proline-rich motif of RhoU N-terminal extension constitutes the preferential binding site for the SH3 domains of ITSN proteins.

**Figure 2:**
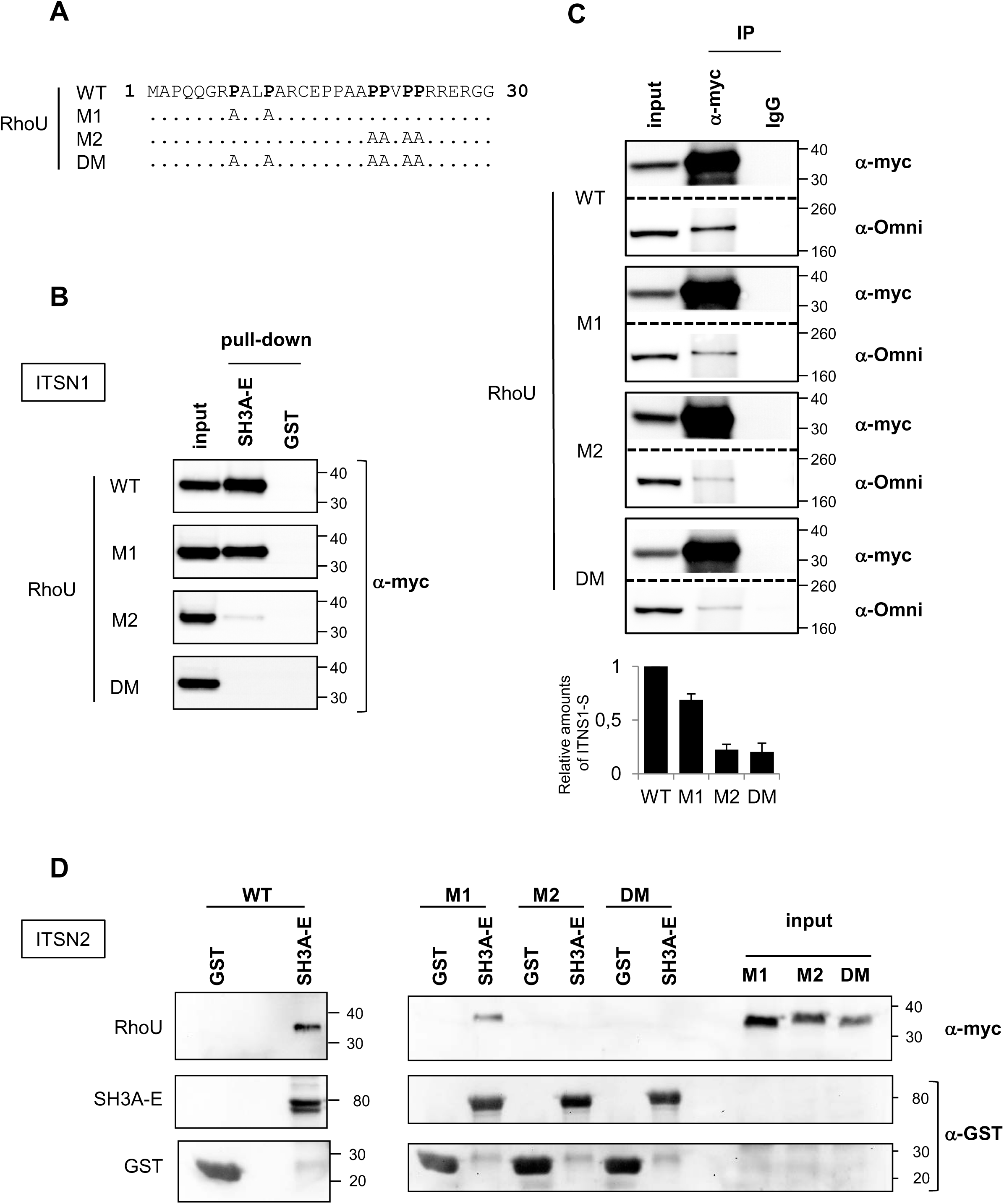
Proline-rich motifs of RhoU mediate the interaction with SH3 domains of ITSNs. **(A)** Schematic representation of RhoU proline-rich mutants used. Proline to alaninesubstitution are indicated. **(B)** Lysates from cells expressing either WT, M1, M2 or DM RhoU mutants were incubated with either GST alone or GST fused to the five SH3 domains of ITSN1 (SH3A-E). Precipitated RhoU was revealed by western blot using anti-myc antibodies. **(C)** Coimmunoprecipitation experiments from lysates of cells expressing myc-tagged RhoU and Omni-tagged ITSN1-L. Myc-tagged RhoU was precipitated using anti-myc antibodies and associated ITSN1 revealed by western blot. Relative amounts of precipitated ITSN1 are shown on the graph. **(D)** GST or GST SH3A-E domains of ITSN2 were used on lysates from cells expressing WT RhoU and indicated RhoU mutants. Precipitated RhoU and recombinant proteins were revealed by western blot using anti-myc and anti GST antibodies respectively.

### ITSN2 and RhoU are involved in transferrin receptor trafficking pathway

ITSN proteins have been mostly involved in clathrin-mediated endocytosis by participating to clathrin coated pits maturation (Henne et al., 2010; Praefcke et al., 2004; Tsyba et al., 2011). Moreover, RhoU has been localized to early endosomes (Zhang et al., 2011), a major sorting platform of endocytic pathways. Together with our *in vitro* observations, this suggests that RhoU and ITSN2 may be involved in the same pathway to regulate endocytosis and/or recycling of cargo engaged into the clathrin-mediated endocytic pathways. To test this hypothesis, we examined whether transferrin (Tf) endocytic and recycling pathways were altered following depletion of ITSN or RhoU proteins by siRNA (Fig.3A). First, using flow cytometry, we evaluated Tf endocytosis by continuously feeding HeLa cells depleted for ITSN1, ITSN2 or RhoU for 15 min at 37°C with Alexa-Fluor-647-labeled Tf (Tf-A647). After stripping off Tf-A647 bound to cell surface, we measured mean fluorescence intensity of endocytosed Tf-A647. We observed a modest but significant increase in Tf-A647 uptake following depletion of ITSN2 or RhoU (Fig.3B). In contrast, depletion of ITSN1 had no effect (Fig.3B). As a control, clathrin heavy chain (CLTC) depletion showed that clathrin-dependent endocytosis was efficiently inhibited in our assays (Fig.3B). To get further insight into the subcellular route altered, we next set up a fluorescent-based image analysis to quantify both levels of Tf-A647 in cells and in vesicles. In agreement with our flow cytometry experiments, HeLa cells fed for 15 min with Tf-A647 showed increased amounts of Tf upon either ITSN2 or RhoU silencing as compared to control or ITSN1 knocked-down cells (Fig.3D). Interestingly, Tf-positive vesicles were more dispersed in cells silenced for ITSN2 or RhoU compared to control cells whereas there was no apparent change in Tf distribution in ITSN1 knocked-down cells (Fig.3C). Since RhoU and ITSN2 proteins interacted and regulated Tf trafficking, we next looked at the subcellular distribution of RhoU and ITSN2 co-expressed in HeLa cells. In agreement with previous studies (Henne et al., 2010; Pucharcos et al., 2000), HA-ITSN2-L localized to small puncta at the cell periphery (arrowheads, inset 1, Fig.3E) reminiscent of clathrin coated pits (CCP). RhoU was found in linear structures at the cell periphery resembling focal adhesions (arrows, inset 1, Fig.3E). Interestingly, neither GFP-RhoU localized to peripheral ITSN2 puncta nor ITNS2 was found at focal adhesions stained for RhoU. However, we did observe colocalization between RhoU and ITSN2 in a subset of perinuclear vesicles (arrowheads, inset 2, Fig.3E). We could also observe ITSN2 enriched in small tubules emanating from GFP-RhoU vesicles (arrows, inset2, Fig.3E). Altogether, these experiments revealed the importance of RhoU and ITSN2 in the regulation of the Tf endocytic pathway.

**Figure 3:**
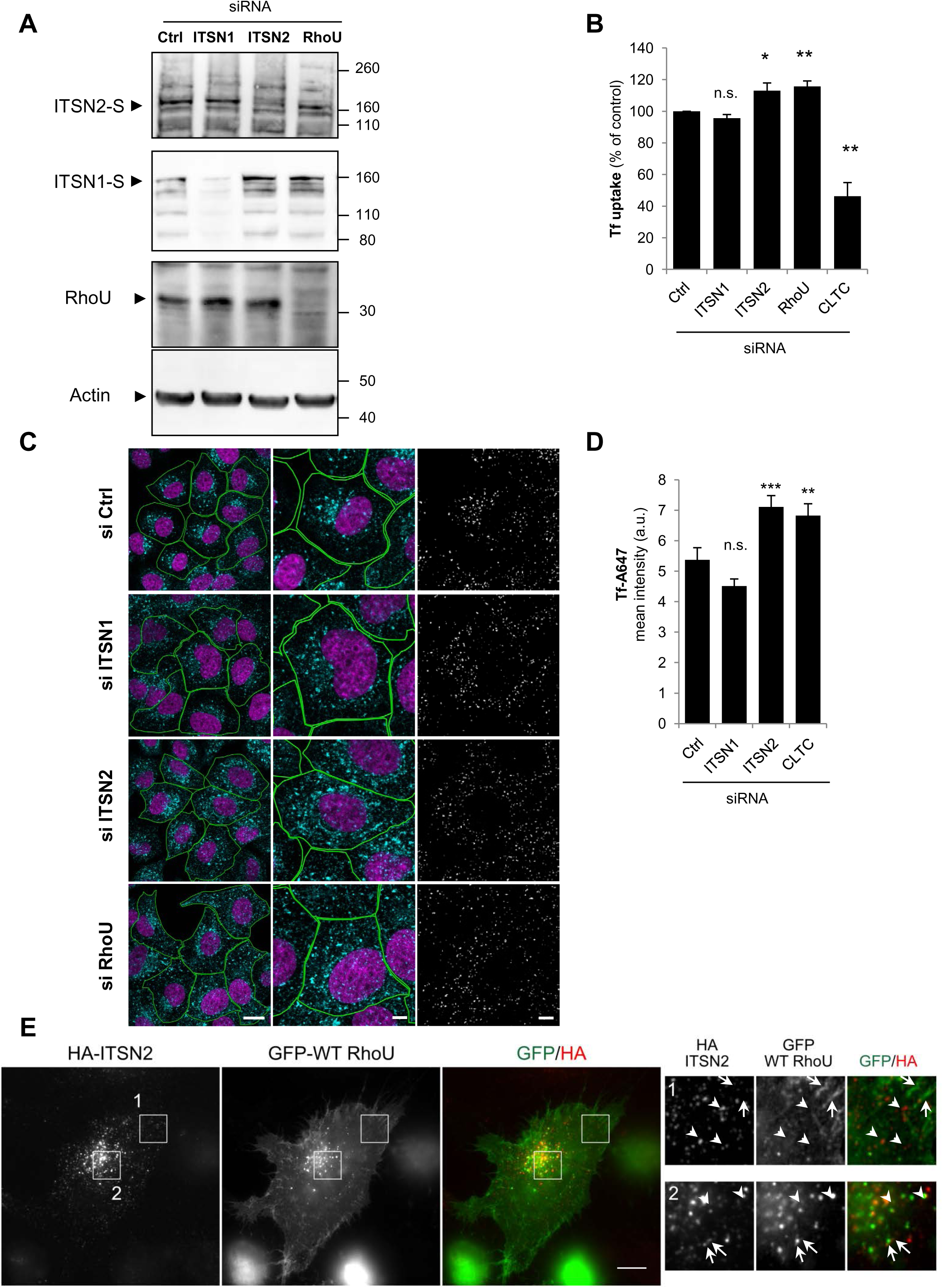
Tf trafficking is defective in HeLa cells silenced for RhoU or ITSN2. **(A)** Western blot analysis of lysates from HeLa cells transfected with siRNA targeting ITSN1, ITSN2 or RhoU compared to cells transfected with an unrelated siRNA (Ctrl). **(B)** Flow cytometry quantification of the endocytosis of transferrin. HeLa cells silenced for ITSN1, ITSN2, RhoU or clathrin heavy chain (CLTC) were serum starved for 30 min prior to incubation with Tf-A647 for 15 minutes. Cells were detached and the mean intensity of internalized Tf in at least 30 000 cells was determined by flow cytometry. Data are represented as percentages of control cells (n=2 experiments). One way ANOVA with Holm-Sidak post-hoc test was used to assess significance *, p<0.05; **, p<0.01; ***, p<0.001 n.s., not significant as compared with control. **(C)** Left: representative field of view of cells silenced for ITSN1, ITSN2 or RhoU and subjected to 15 minutes fluorescent Tf uptake experiments. Nucleus (magenta) and Tf (cyan) are shown. Bar: 10 μm. Middle: a close up view of single cell is shown. Bar: 5 μm. Right: binary mask corresponding to segmented fluorescent spots of Tf-positive vesicles using wavelet spot detector in Icy software. Bar: 5 μm. **(D)** Quantification of Tf endocytosis. The sum of vesicle intensity was computed and normalized to the cell surface and the mean intensity for each condition shown on the graph. One way ANOVA with Holm-Sidak post-hoc test was used to assess significance. **, p<0.01; ***, p<0.001; n.s., not significant as compared with control. **(E)** Wide field immunofluorescence of HeLa cells transfected with HA-ITSN2-L and myc-tagged WT RhoU. Inset1 shows peripheral localization of ITSN2 in puncta (arrowheads) and GFP-RhoU in focal adhesions (arrows). Inset2 shows partial colocalization of ITSN2 and GFP-RhoU in perinuclear vesicles. Arrow and arrowheads point to vesicles with both ITSN2 and GFP-RhoU. Note that ITSN2 can be found enriched at tips of tubules emanating from GFP-RhoU vesicle (arrows). Bar 5 μm

### ITSN2 and RhoU regulate transferrin receptor recycling

We next attempted to unravel how RhoU and ITSN2 together regulate Tf receptor trafficking. The above increase in intracellular Tf can result from more receptor at the cell surface and/or a delay in Tf recycling leading to accumulation of Tf in cells. To estimate the amounts of Tf receptor at the cell surface, we incubated cells with Tf-A647 for 1 hour at 4°C, a condition that blocks endocytosis. After a quick wash, amounts of bound Tf-A647 were estimated by flow cytometry (Fig.4A). Fluorescence levels were not significantly different in HeLa cells depleted for ITSN2, RhoU or CLTC as compared to the control, indicating that the higher uptake of Tf is probably not the consequence of an increase in cell surface Tf receptor. To test for a defect in Tf recycling, the surface pool of Tf receptor was labeled at 4°C with Tf-A647 for 1 hour. Excess of Tf-A647 was then removed and HeLa cells transferred to 37°C for 5, 15 or 30 minutes. After stripping off the surface pool of Tf-A647, amounts of intracellular Tf-A647 were estimated by flow cytometry. Interestingly, as early as 5 minutes after triggering endocytosis twice the amount of Tf was endocytosed in cells depleted for ITSN2 or RhoU as compared to control cells (Fig.4B). Larger amounts of endocytosed Tf were maintained at 15 and 30 minutes time points but differences were reduced suggesting that the major trafficking defect occurs early during transferrin receptor trafficking process.

**Figure 4:**
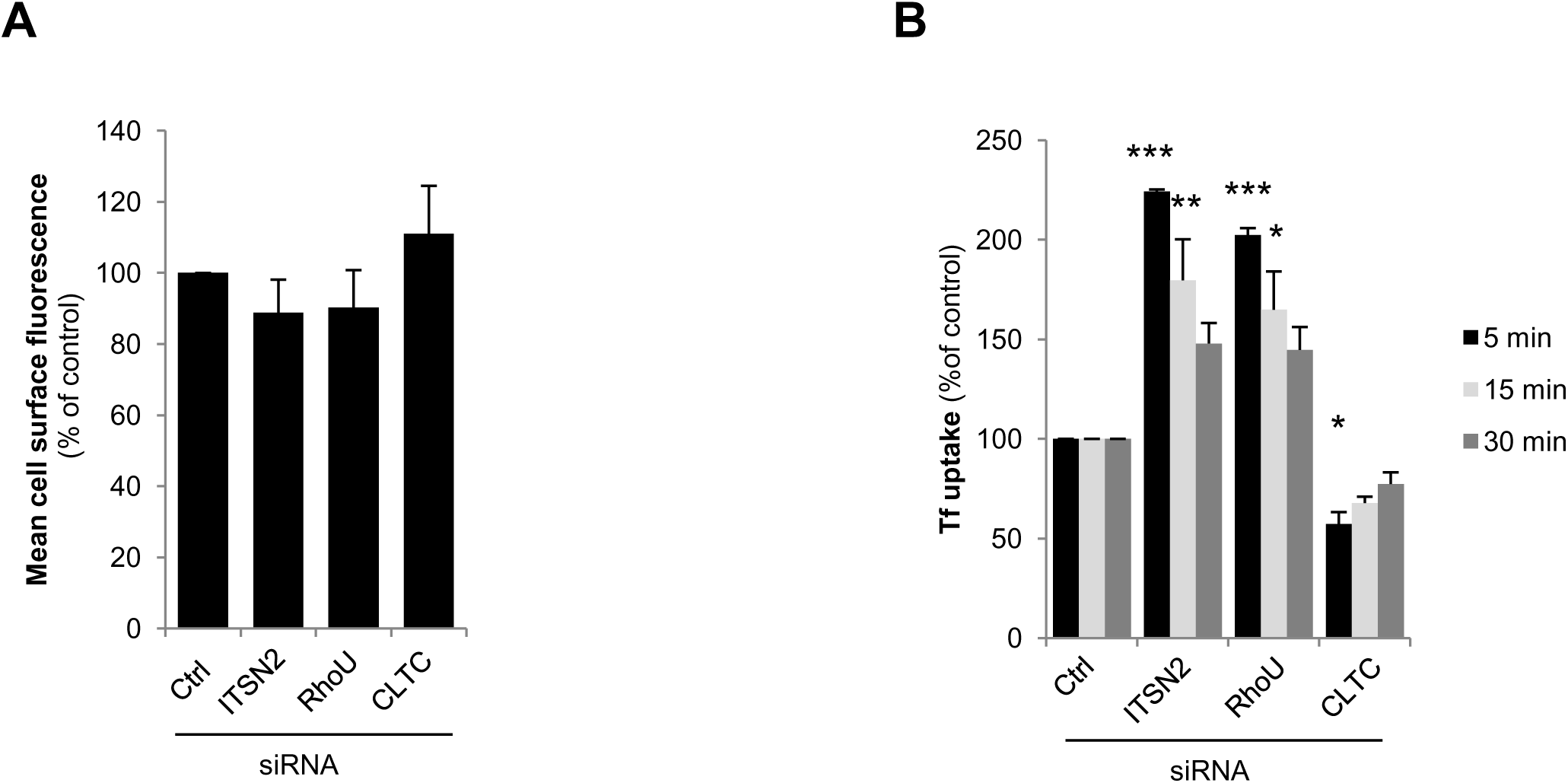
Depletion of ITSN2 or RhoU alters the fast recycling pathway. Hela cells were silenced by siRNA transfection for ITSN1, ITSN2, RhoU or clathrin heavy chain (CLTC) and compared to cells expressing unrelated siRNA (Ctrl). (A) Flow cytometry analysis of Tf-A647 binding at the cell surface. (B) Time course of Tf-A647 endocytosis. Cells were serum starved for 30 min and incubated on ice for 1h with Tf-A647. Cells were quickly washed and incubated for 5, 15 or 30 min at 37°C prior to flow cytometry analysis. The mean intensity of internalized Tf in at least 30 000 cells was determined and data are represented as percentages of control cells considered as 100% (n=2 experiments). One way ANOVA with Holm-Sidak post-hoc test was used to assess significance. ***, p<0.001; **, p<0.01; *, p<0.05.

Once endocytosed, Tf receptors can be recycled back from endosomes to the plasma membrane by two recycling modes, “a fast” one occurring within 5 minutes directly from early endosomes or “a slow” one after sorting to recycling endosomes occurring after 15 min (Ciechanover et al., 1983; Grant and Donaldson, 2009). Therefore, one possible explanation for the early transferrin increase in response to ITSN2 or RhoU silencing is an alteration of the fast recycling mode. To test this hypothesis, we performed pulse chase experiments to follow the distribution of Tf-A647 in HeLa cells by immunofluorescence (Fig.5). Since the first steps of endocytosis requires roughly 2 minutes before fast recycling takes place (Aguet et al., 2013; Hao and Maxfield, 2000; Taylor et al., 2011), we labeled surface Tf receptor with Tf-A647 for 1 hour at 4°C, then placed cells at 37°C for 2 minutes to stay within the limit of “pure endocytosis”. After stripping off cell surface Tf-A647 at 4°C, cells were either fixed (t=0 in Fig.5) or returned back to 37°C for 2 or 5 minutes before fixation, to analyze the fate of Tf-A647 that had been endocytosed during the 2 minutes pulse. We then quantified two parameters: first, we calculated the percent of Tf-positive vesicles that colocalized with EEA1 compartments (object based colocalization) and second, the percentage of Tf-A647 found in EEA1 compartments at a given time by integrating amounts of fluorescent Tf in EEA1 compared to total amounts of Tf in cells. As shown in Fig.5A, in control cells, a small proportion of Tf-positive vesicles had already reached EEA1 compartment at t=0. As previously reported (Kalaidzidis et al., 2015), colocalization of Tf-containing vesicles with EEA1 increased between 0 and 2 min and started to decrease between 2 to 5 min suggesting that either Tf had been recycled or had left the EEA1 compartment to reach the recycling endosomes. In cells depleted for ITSN2 or RhoU, amounts of Tf-containing vesicles colocalizing with EEA1 was comparable to control cells (Fig.5A,B) indicating that, at early time point, there is no major rerouting of Tf. However, when we estimated amounts of Tf in EEA1 compartment, we found that, at 2 and 5 minutes, amounts of fluorescent Tf in EEA1 was increased after depletion of ITSN2 or RhoU (Fig.5C) suggesting that Tf residency time in EEA1 was increased in these conditions. To further analyze the fate of Tf-A647, we conducted pulse-chase experiments for 2 and 5 minutes and estimated amounts of Tf-A647 reaching Rab11-positive endosomes. Fig.5D shows that in agreement with a delay in recycling or a slowing down of vesicle maturation toward recycling endosomes, the proportion of Tf in Rab11 compartment was higher at 2 and 15 min when ITSN2 or RhoU was silenced. Altogether, these data demonstrate that RhoU and ITSN2 regulate Tf-vesicle progression along the endosytic route and might be needed for fast recycling of Tf receptor.

**Figure 5:**
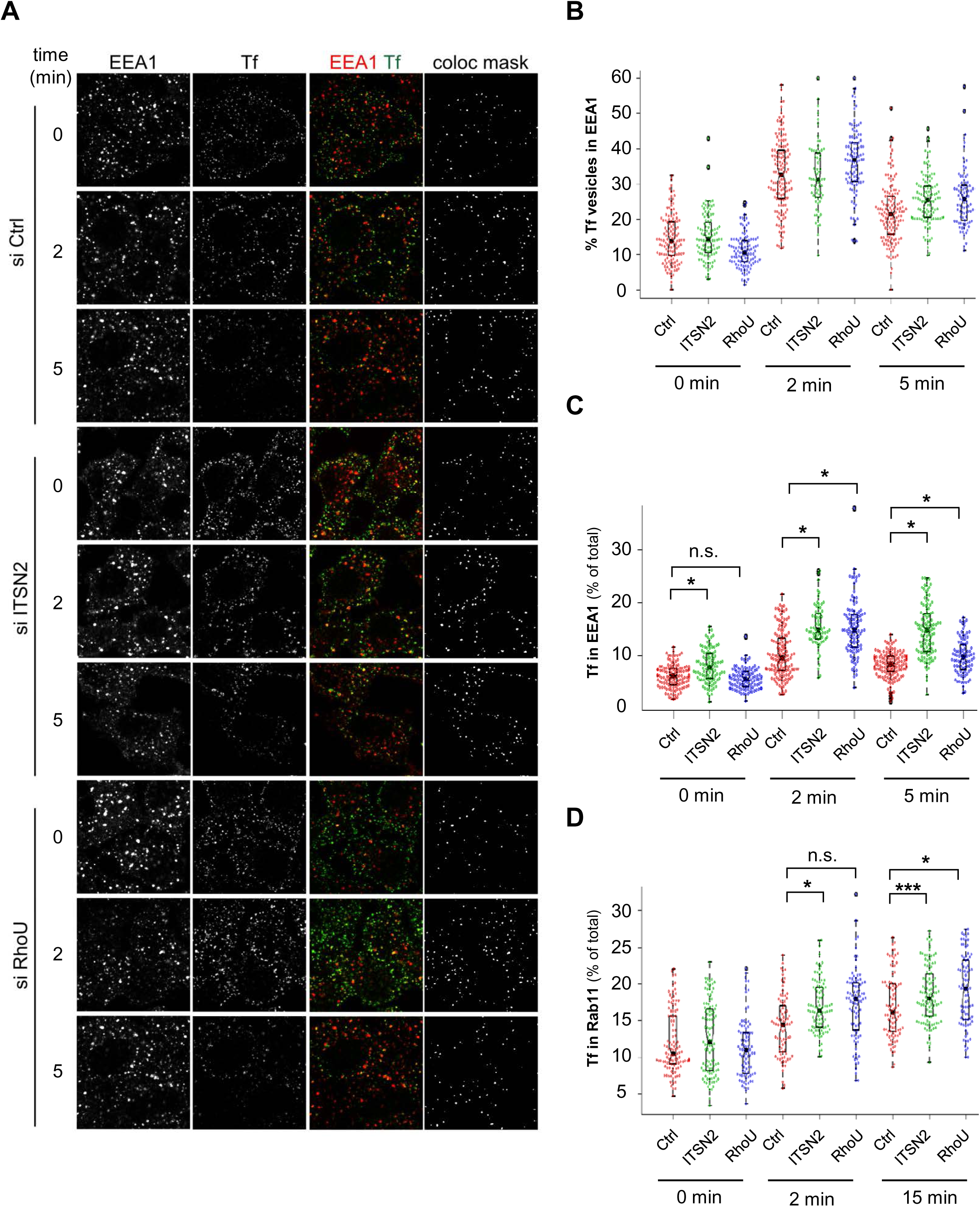
Depletion of ITSN2 or RhoU leads to accumulation of Tf into EEA1 compartments. **(A)** Representative field of view of control HeLa cells (si Ctrl) and Hela cells silenced for ITSN2 or RhoU pulsed for 2 min with Tf-A647 and chased for indicated time after stripping off cell surface Tf. Cells were stained for EEA1 and the colocalization mask between EEA1 and Tf-A647 shown. The percentage of Tf-positive vesicle colocalizing with EEA1**(B)**, the fraction of Tf-A647 found in EEA1- **(C)** or Rab11-positive endosomes **(D)** as compared to total amounts of Tf in cells were quantified overtime (n=2 experiments, at least 40 cells/experiments and conditions). To assess significance, Kruskal-Wallis on rank with Dunn’s post-hoc test was performed for (B) and (C) and one way ANOVA with Holm-Sidak post-hoc test for (D). ***, p<0.001; *, p<0.05; n.s.: not significant)

### RhoU is distributed along the endocytic route

RhoU has been localized into cell adhesion structures and vesicular compartments (Chuang et al., 2007; Ory et al., 2007; Zhang et al., 2011). The precise distribution of RhoU along the endocytic route is however unknown. Since available antibodies against RhoU do not detect endogenous RhoU by immunofluorescence, we co-transfected myc-RhoU together with GFP-tagged Rab4a, 5a, 11a and 7a, several specific markers of the endocytic route (Wandinger-Ness and Zerial, 2014). RhoU was found partially colocalized with fast recycling endosomes (Rab4a), early endosomes (Rab5a) and slow recycling endosomes (Rab11a; Fig.6A). Very weak colocalization was observed with late endosomes. Indeed, less than 10% of WT-RhoU was found in GFP-Rab7a compartments (not shown) as compared to around 30% of RhoU colocalized with GFP-Rab4a and GFP-Rab5a and 50% with GFP-Rab11a (Fig.6B). Since DM RhoU is impaired in ITSNs binding, we used this mutant to test if the subcellular distribution of RhoU depends on its ability to interact with ITSNs. Fig.6B showed that the distribution of DM RhoU was comparable to WT RhoU suggesting that ITSNs binding is dispensable for RhoU subcellular distribution.

**Figure 6:**
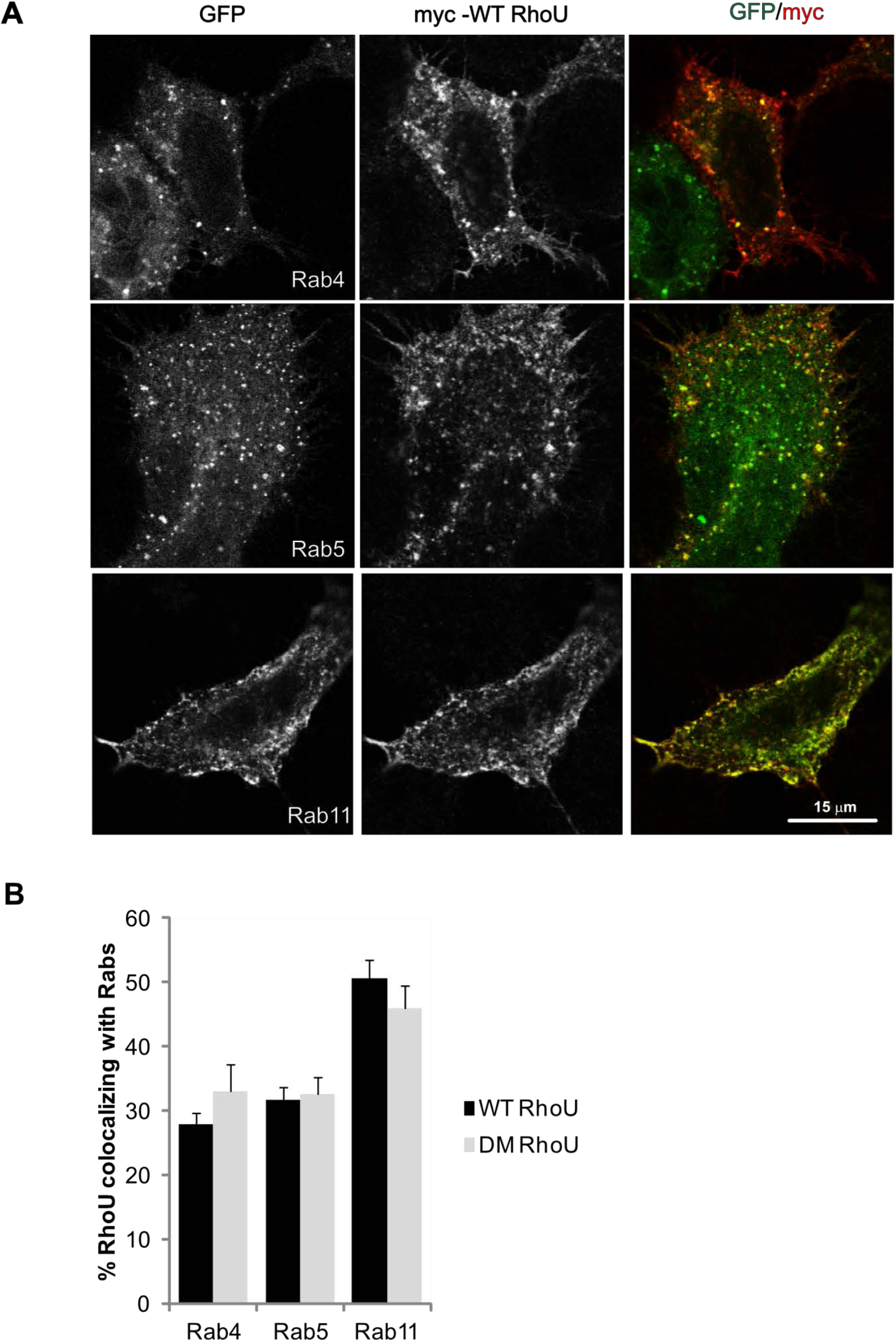
RhoU is distributed all along the endocytic route. **(A)** Confocal microscopy analysis of HeLa cells co-expressing GFP Rab4a, GFP-Rab5a or GFP-Rab11a and myc-tagged WT RhoU. **(B)** Quantification of RhoU colocalizing with Rab proteins. HeLa cells were transfected with GFP-Rab proteins along with myc-WT RhoU or myc DM-RhoU and the percent of colocalization determined. Graph represents mean of percent +/- sem (n=2 experiments, 15 cells/experiments).

### ITNS2 partially colocalized with RhoU on Rab4 compartment

Since delays in Tf recycling occurred early after the pulse, we sought for colocalization of transfected myc-RhoU and HA-ITSN2 (S and L forms) with GFP-Rab4a, a marker of fast recycling endosomes (Grant and Donaldson, 2009). Although amounts of colocalization of HA-ITSN2, myc-RhoU and GFP-Rab4a were low, we could find few vesicular structures containing RhoU and Rab4a with discrete amounts of ITSN2L (Fig.7A) or ITSN2S (Fig.7B). This suggests that ITSN2 could transiently associate with early endosomes that may contain Rab4a and RhoU (Sönnichsen et al., 2000; Zhang et al., 2011). To further confirm that ITSN2 could associate with early/sorting endosomes, we used spinning disk confocal microscopy to perform live cell imaging of HeLa cells co-transfected with GFP-ITSN2S and tagRFP-T-EEA1, a fluorescent tagged EEA1 that colocalizes with the endogenous proteins (Kalaidzidis et al., 2015; Navaroli et al., 2012). This revealed that there were in fact vesicles containing both EEA1 and ITSN2 (Fig.8A). These vesicles are motile and can move toward an immobile EEA1 vesicle. Once in close contact, both types of vesicle remained associated for roughly 1 minute to eventually segregate into two distinct vesicles, one containing both EEA1 and GFP-ITSN2S and another one containing only EEA1 (Fig.8A,B). Interestingly, vesicle motility analysis indicated that a fraction of EEA1 and ITSN2 vesicles were motile (Fig.8C) and they can associate for a short period of time (Fig.8D). Indeed, live cell colocalization analysis showed that 15% of total vesicles were double stained for EEA1- and ITSN2 for less than 20 seconds. These observations suggest that subset of EEA1- and ITSN2-vesicles transiently associated to potentially regulate the fate of cargo.

**Figure 7:**
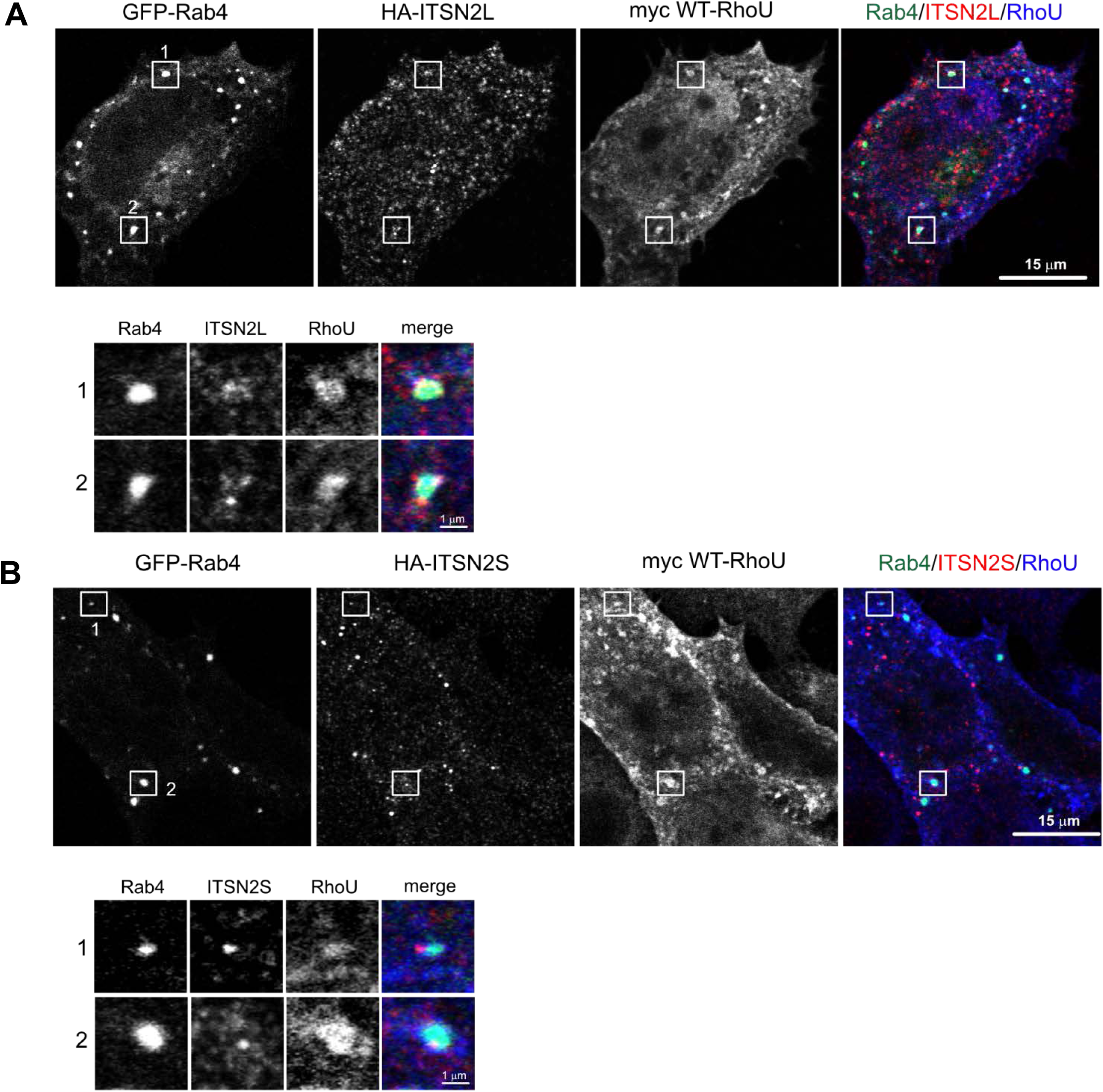
ITSN2 and RhoU colocalized to Rab4-positive endosomes. **(A)** Confocal microscopy analysis of Hela cells transfected with GFP-Rab4, HA-ITSN2-L and myc-WT RhoU. Boxes show regions magnified in inset 1 and 2. **(B)** Confocal microscopy analysis of Hela cells transfected with GFP-Rab4, HA-ITSN2-S and myc-WT RhoU. Boxes show regions magnified in inset 1 and 2.

**Figure 8:**
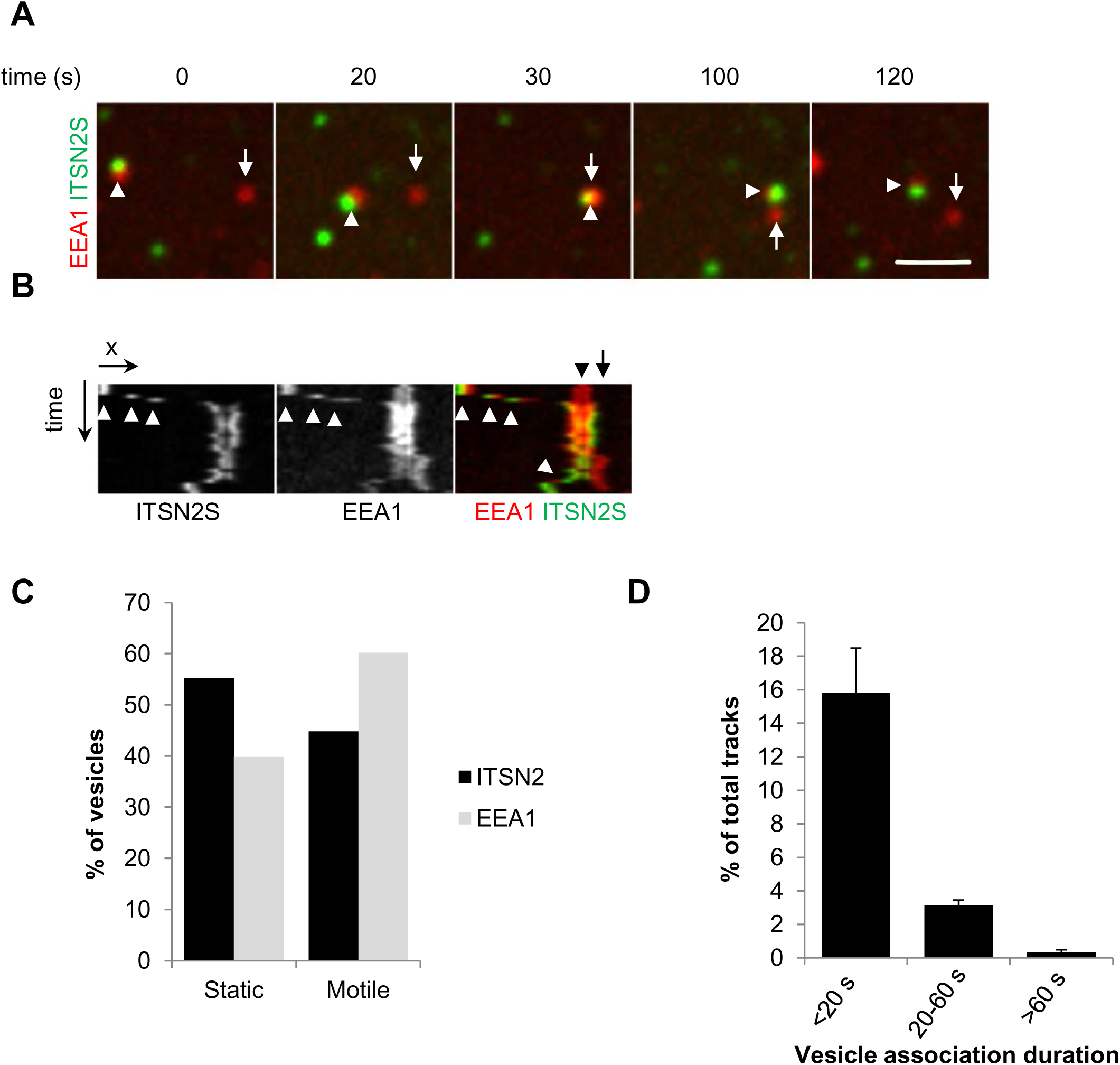
ITSN2-S is transported with EEA1-positive endosomes. **(A)** Still images of HeLa cells co-transfected with GFP-ITSN2-S and TagRFP-T-EEA1. Cells were imaged with spinning-disk confocal microscope for 2 min at 0.5 Hz. Arrowhead points to endosome containing both ITSN2-S and EEA1. Arrow points to single stained EEA1-positive endosome. **(B)** Kymograph of the sequence. White arrowheads points to endosomes co-satined with ITSN2-S and EEA1 and black arrowhead to single stained EEA1 endosome. Double stained move toward EEA1 endosome and eventually segregate (white arrowhead, double stained; black arrow, EEA1-positive only). **(C)** Quantification of static and motile vesicles from the movie. 144 ITSN2- and 107 EEA1-positive vesicles were detected. Vesicles with displacement below 0.7 μm were considered static. **(D)** Quantification of ITSN2- and EEA1-positive vesicles that remained associated over time. ITSN2 and EEA1 vesicles were tracked overtime and considered associated if tracks of vesicle centroids were separated by 3 pixels or less. Associations lasting less than 6 seconds (3 consecutive images) were discarded for the analysis. Tracks corresponding to double stained vesicles were counted and normalized to the total number of tracks (sum of ITSN2 and EEA1 tracks) and expressed as percent +/- sem (n=4 cells).

## Discussion

RhoU is an atypical Rho GTPase mostly involved in the regulation of cell migration and transformation (Berzat et al., 2005; Brady et al., 2009; Brazier et al., 2009; Chuang et al., 2007; Fort et al., 2011; Ory et al., 2007; Zhang et al., 2011). RhoU has been localized to diverse subcellular compartments including focal adhesions, the plasma membrane and intracellular vesicles. Although RhoU associates to endosomal compartments (Alan et al., 2010; Shutes et al., 2004; Zhang et al., 2011), no studies have addressed the potential function of RhoU in vesicle trafficking. Here, by looking for new RhoU binding partner and further characterizing the subcellular distribution of RhoU along the endocytic route, we provide evidences that RhoU binding to ITSN2 may regulate fast recycling of endocytosed cargo. Indeed, we identified ITSNs as part of RhoU complex and showed that the second PxxPxR motif of RhoU acts as the preferential docking interface between RhoU and SH3 domains of ITSNs. Moreover, we found that silencing of ITSN2 but not ITSN1 reduced Tf recycling leading to a global increase of Tf uptake. Interestingly, defect in Tf recycling was phenocopied by RhoU silencing indicating that they act in the same pathway. Uptake of transferrin following a short pulse revealed that RhoU and ITSN2 are required for fast recycling of Tf. Concomitantly, immunofluorescence analysis of ITSN2 and RhoU indicated that both proteins partially colocalized with Rab4, a marker of fast recycling vesicles. We therefore propose that RhoU and ITSN2 interaction may occur on a subset of vesicles to drive fast recycling of cargos.

Due to their association with key components of clathrin mediated endocytosis including dynamin, synaptojanin and AP2 complex (Praefcke et al., 2004; Yamabhai et al., 1998), most studies addressing ITSN1 and ITSN2 function have focused on early phase of endocytosis. ITSN1 and ITSN2 are needed to progress from the formation of early nucleation module to the clathrin-coated pit (CCP) by bridging FCHo and AP-2 (Henne et al., 2010; Praefcke et al., 2004). In contrast to what we had anticipated, the silencing of ITSN1 or ITSN2 did not inhibit Tf endocytosis but instead, either increased Tf accumulation in cells silenced for ITSN2 or had no effect in the case of ITSN1 silencing. Although surprising, our results are nonetheless consistent with previous reports showing that ITSNs silencing have limited effect on Tf endocytosis (Das et al., 2007; Martin et al., 2006; Russo and O’Bryan, 2012; Yang et al., 2015) and with the idea that ITSN1 and ITSN2 can compensate each other in early phase of clathrin coated pit assembly (Henne et al., 2010). The fact that Tf accumulation was maximal after a short pulse of Tf uptake in the absence of RhoU or ITSN2 indicated that recycling defects occurred shortly after completion of CCP assembly. Accordingly, we detected accumulation of Tf in EEA1-positive endosomes which constitute the main sorting platform of endocytosed cargos. Endosomes can be seen as a dynamic arrangement of membrane domains containing various amounts of Rab proteins that regulate the fate of vesicle (Rink et al., 2005; Sönnichsen et al., 2000) and our colocalization analysis revealed that RhoU colocalized with Rab5, Rab4 and Rab11 but much less with Rab7 indicating that RhoU is most likely involved in recycling of cargo rather than its degradation. Since RhoU colocalized with ITSN2 on a subset of Rab4 containing vesicles, a marker of fast vesicle recycling, it suggests that the RhoU-ITSN2 interaction may constitute a molecular switch to drive fast recycling of vesicular cargos. The use of fluorescent Tf as a readout of endocytosis does not exclude that other cargo including receptors or adhesion molecules might be regulated by RhoU-ITSN2 interaction. Indeed, despite various mode of entry, cargos eventually meet to EEA1-positive endosomes from where they will be recycled back to the plasma membrane or degraded (Barysch et al., 2009; Lakadamyali et al., 2006; Leonard et al., 2008). Interestingly, RhoU has been involved in several biological processes that may depend on plasma membrane protein recycling like, for example, cell migration (Chuang et al., 2007; Fort et al., 2011; Ory et al., 2007) and cell polarity (Brady et al., 2009). Whether RhoU and ITSN2 control cell adhesion molecule turnover has to be investigated but Rab4-dependent recycling of integrins at the plasma membrane is key to cell motility and cell adhesion turnover (Arjonen et al., 2012; Roberts et al., 2001) and both RhoU and ITSN2 silencing alters cell-cell junction and epithelial cyst formation in a three-dimensional matrix (Brady et al., 2009; Qin et al., 2010). RhoU has also been linked to EGFR signaling thanks to Grb2 binding. Upon EGF stimulation, endosomal RhoU is recruited by Grb2 in EGFR complex to sustain JNK activation. Grb2 binds to RhoU through the same motif as ITSN2 (Zhang et al., 2011). Whether ITSN2 and Grb2 may compete for RhoU binding and control the fate of the EGFR would be an interesting issue to address. Finally, RhoU expression levels are regulated by various signals including Wnt-1, RANKL and Notch to induce cell differentiation (Bhavsar et al., 2012; Brazier et al., 2006; Schiavone et al., 2009; Tao et al., 2001). Therefore, according to its expression levels, RhoU might modulate the rate of endocytosis and/or recycling of receptors to control signal output and cell fate.

The subcellular localization analysis of RhoU and ITSN2 indicated that both proteins are mutually excluded from plasma membrane compartments where RhoU or ITSN2 accumulates. RhoU is found in focal adhesions but not in CCP where ITSN2 is enriched. Conversely, ITSN2 is not localized to focal adhesions indicating that RhoU and ITSN2 may achieve distinct functions and meet on endosomes to ensure specific cargo recycling. Vesicular localization of ITSNs suggest also that, in addition to their role in CCP maturation, they may have a function in vesicle trafficking. Few reports suggest indeed that ITSNs might regulate endosomal trafficking. For example, yeast two-hybrid screen have identified Rabaptin-5 and the kinesin KIF16B as potential binding partners of ITSN1 and ITSN2 (Wong et al., 2012; Yang et al., 2015). Both Rabaptin-5 and KIF16B coordinate Rab5 and Rab4 for efficient vesicle recycling (Hoepfner et al., 2005; Pagano et al., 2004; Stenmark et al., 1995). Recently, ITSN1-S has been reported to localize to Rab4-positive vesicles (Gryaznova et al., 2018) and to interact with DENND2B, an exchange factor for Rab13 involved in EGF receptor recycling (Ioannou et al., 2017). ITSN1 knock-out mice showed only minor (Yu et al., 2008) or no defects (Sakaba et al., 2013) in CME-dependent synaptic vesicle recycling in neurons, but neurons have enlarged EEA1-endosomes (Yu et al., 2008). Together with our data showing that ITSN2 localizes to Rab4 and associates to EEA1-moving vesicle in living cells, it provides clues for potential roles of ITSNs in vesicle transport and recycling. The fact that ITSN1 and ITSN2 silencing had differential consequence on Tf recycling suggests that if they can compensate each other in CCP maturation, they may have distinct functions in vesicular trafficking.

Our findings question whether RhoU activities might be regulated by ITSNs and/or RhoU might modulate Cdc42 activities. RhoU is believed to be constitutively active since it is GDP/GTP exchange rate *in vitro* is spontaneous and fast. However, the deletion of its N-terminus containing PxxP motifs enhances its ability to bind and activate PAK to promote cell transformation suggesting that RhoU N-terminus negatively regulates its activity (Shutes et al., 2004). Modulation of RhoU activities have been proposed to occur in response to EGF signaling in which binding of Grb2 to RhoU may relieve N-terminus dependent inhibition (Zhang et al., 2011). Therefore, ITSNs binding to RhoU might be a way to locally enhance RhoU activities. Further work will be needed to develop tools able to monitor local RhoU activities. Remarkably, by testing individual SH3 domains, we found that 3 domains of ITSN1 (SH3A, C, E) and only one domain of ITSN2 (SH3E) was mediating RhoU binding. This discrimination is rather surprising considering that SH3A, C and E domains of ITSNs fall into canonical classes of SH3 domains and preferentially binds to classII PxxPxR motifs (Teyra et al., 2017). It also suggests that depending on the ITSN engaged with RhoU, the stoichiometry of the ITSN/RhoU complex might be different.

Finally, ITSNs are well known exchange factors for Cdc42 thanks to their Dbl homology (DH) domain. The long form of ITSNs is less potent to activate Cdc42 than the DH domain alone suggesting that SH3 domains of ITSN negatively regulates DH-dependent nucleotide exchange activity (Zamanian and Kelly, 2003). RhoU binding to ITSNs may therefore relieve auto inhibition and facilitate local Cdc42 activation. Activated Cdc42 controls filopodia formation and cell polarity, two features shared by RhoU (Alan et al., 2010; Brady et al., 2009; Ruusala and Aspenström, 2008). By binding to WASP proteins, ITSN2-L facilitates Cdc42-dependent actin polymerization (Klein et al., 2009; McGavin et al., 2001). Actin polymerization may act as a scaffold to limit cargo diffusion on endosomes and control their fate (Simonetti and Cullen, 2019). In contrast to Cdc42, RhoU does not interact with the GTPase binding domain of WASP (Saras et al., 2004) and among RhoU effector, none has been reported to mediate actin polymerization. Our study therefore raises new issues on a potential cross-talk between RhoU, ITSN2 and Cdc42 to regulate sorting of cargoes.

The authors declare no competing financial interests

## Acknowledgements

We acknowledge the generosity of Dr Susana de la Luna (CRG, Barcelona, Spain), Dr Daniel Billadeau (Mayo Clinic, Rochester, MN, USA) and Dr Philippe Chavrier for kindly providing us with human ITSN2, mouse RhoU and human Rabs coding plasmids respectively. We acknowledge the confocal microscopy facilities of Plateforme Imagerie In Vitro, the cytometry facility at INCI and Dr Jean-Daniel Fauny (IBMC, UPR3512, Strasbourg) for technical assistance with spinning disk confocal microscopy.

## Materials and Methods

### Constructs and antibodies

The human RhoU was PCR amplified as a BamHI/EcoRI fragment from pRK5-myc constructs (a kind gift from Dr Aspenström, Uppsala, Sweden) and subcloned into the pGEX-4T1 between the BamHI/EcoRI sites.

The constructs encoding the myc-tagged mouse RhoU (WT or proline to alanine mutants) were a kind gift from Dr Billadeau (Rochester, MN, USA) and were described previously (Zhang et al., 2011). The constructs encoding SH3 domains of ITSN1 or ITSN2 fused to GST and Omni-tagged ITSN1L were as described previously (Gryaznova et al., 2015; Tsyba et al., 2008). The cDNAs encoding the human ITSN2-L were a kind gift from Dr S. de la Luna (Barcelona, Spain). The full coding sequence of ITSN2-L was subcloned into the BamHI site of the pEGFP-C1 plasmid. The plasmids encoding GFP-tagged human Rab proteins (Rab4a, Rab5a, Rab11a, Rab7a) were a king gift from Dr P. Chavrier (Institut Curie, Paris, France). The rabbit polyclonal anti-EEA1 was purchased from Santa Cruz, the monoclonal anti-myc (clone 4A6) and the polyclonal anti-RhoU antibodies were from Millipore (Saint Quentin Fallavier, France), the polyclonal anti ITSN2 were from Novus Biologicals (Lille, France), the polyclonal anti-Rab11 from ThermoFisher Scientific (Courtaboeuf, France). Unlabelled and fluorescent transferrin were from Sigma (Saint Quentin Fallavier, France). Mouse polyclonal rabbit α-Omni (M-19) antibodies were from Santa Cruz Biotechnology (USA). Rabbit polyclonal α-GST antibodies were from CliniSciences (France).

### Cell culture and transfection

HEK293T and HeLa cells were cultured in Dulbecco’s modified Eagle’s medium (DMEM) supplemented with 10% fetal bovine serum at 37°C in 5% CO_2_. Plamids DNA were transfected according to manufacturer’s instruction using jetPEI^TM^ (Polyplus, Illkirch, France).

For siRNA transfection, 2×10^4^ HeLa cells/cm^2^ were seeded 24 h prior to siRNA transfection according to manufacturer’s instructions. Lipofectamine RNAiMax (ThermoFisher Scientific, Courtaboeuf, France) and 50 nM of a mix of 4 siRNA directed against *ITSN1, ITSN2, CLTC or RhoU* (On Target Plus Smart Pool siRNA; Dharmacon, Cambridge, UK) were used (see Table 1 for sequences). Cells were cultured for 48 h before experiments and silencing was estimated and normalized to actin contents by western blotting. We obtained more than 70% silencing for each protein in each experiment.

**Table 1:**
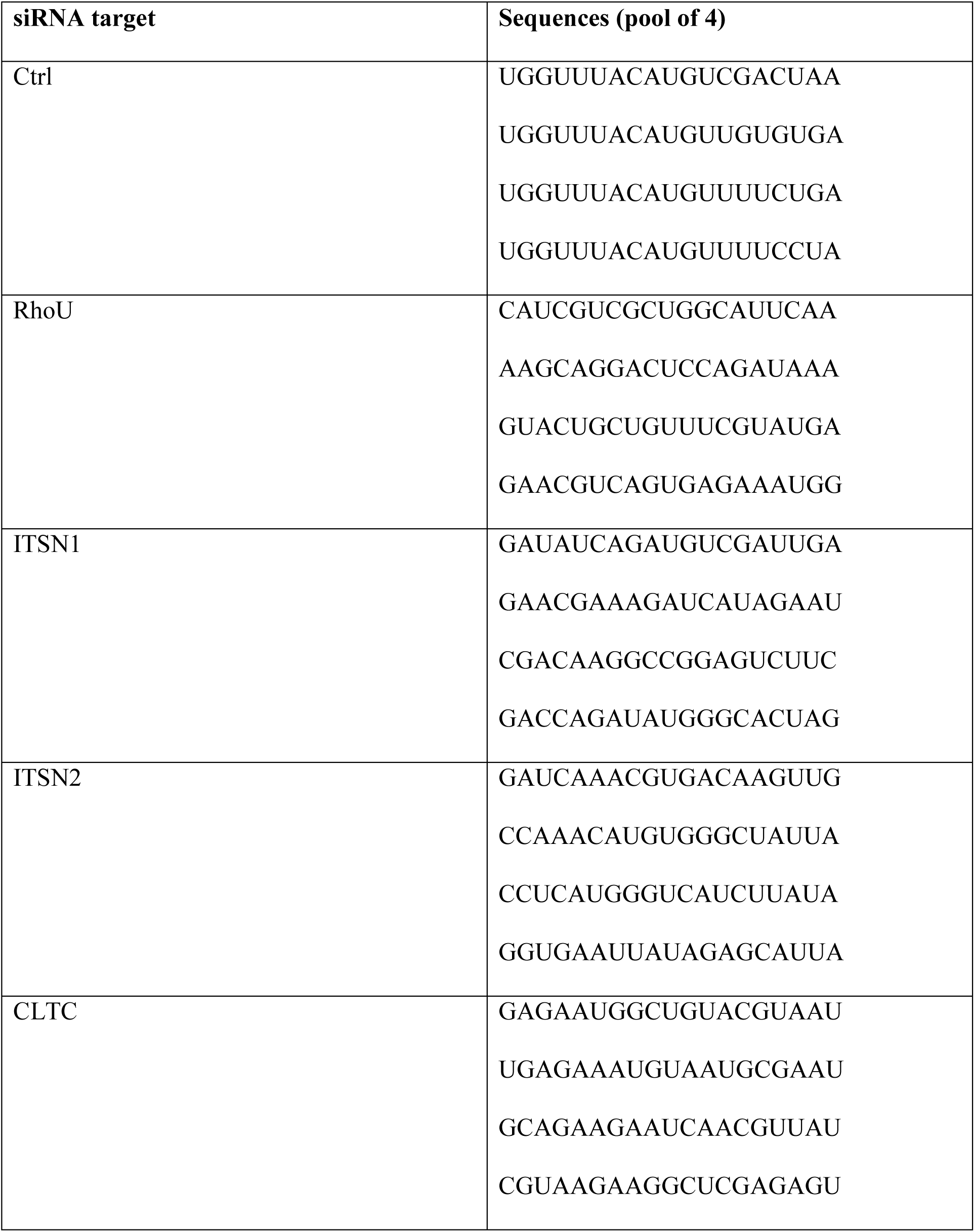
Sequences of siRNA used for this study

### GST Pull down and coimmunoprecipitation assays

GST fusion proteins were produced in *Escherichia coli* BL21 cells. After isopropylthiogalactoside (IPTG) induction, pellets of bacteria were resuspended in lysis buffer containing 50 mM Tris-HCl (pH 8), 2 mM MgCl_2_, 0.2 mM Na_2_S_2_O_5_, 10% glycerol, 20% sucrose, 2 mM DTT, protein inhibitor cocktail (Roche, Mannheim, Germany) and then sonicated. Cell lysates were centrifuged 20 minutes at 4°C at 45000 ***g*** and the supernatants were incubated with glutathione-coupled Sepharose 4B beads (Sigma) for 30 minutes at 4°C. After 3 washes with lysis buffer, amounts of GST fusion proteins bound to the beads were estimated using Coomassie stained SDS gels.

HEK293T cells transfected with myc-tagged RhoU (WT or proline mutants) were rapidly washed in ice cold PBS and lysed on ice with 20 mM Tris-HCl (pH 8), 2 mM MgCl_2_, 1% IGEPAL CA-630, 10 μM GDP, 150 mM NaCl and protein inhibitor cocktail. Lysates were cleared for 5 minutes at 17000 ***g*** at 4°C and aliquots were taken from the supernatant to determine the protein concentration in the total cell lysate. 20 μg of bacterially produced SH3 domains of ITSN1 or ITSN2 fused to the GST protein and bound to glutathione-coupled Sepharose beads were added to cell lysate and maintained for 2 hours at 4°C. Beads with bound proteins were washed 4 times in wash buffer (20 mM Tris-HCl pH 8, 150 mM NaCl, 0.1% IGEPAL CA-630, 2 mM MgCl_2_, 10 μM GDP), eluted in Laemmli sample buffer and then analysed by western blot for precipitated RhoU.

For immunoprecipitations, HEK293T cells were lysed in IP buffer (150 mM NaCl, 20 mM Tris, pH7.5, 10% glycerol, 0,5% NP40, 5 mM MgCl_2_, protease inhibitors cocktail (Sigma) and centrifuged for 15 min at 16,000g. Supernatant (2 mg of proteins) was incubated with 2 μg of target antibodies and 20 μl of Protein A/G Plus Ultralink Resin (Thermo Scientific) for 4 hours at +4°C. Then beads were washed four times with IP buffer. Immunoprecipitated complexes were eluted with Laemmli buffer and analyzed by western blot

### Endocytosis, recycling assays and immunofluorescence

For continuous endocytosis assays, HeLa cells were serum starved for 30 minutes in DMEM supplemented with 0.5% BSA (DMEM/BSA) and incubated for 15 minutes with DMEM/BSA containing 10 μg/ml Alexa 647 conjugated transferrin (ThermoFischer Scientific) at 37°C. Cells were transferred on ice and washed twice with ice-cold PBS before fixation for 10 min with 4% (w/v) paraformaldehyde in PBS. Nuclei were stained with Hoechst.

For fast recycling assays, HeLa cells were serum starved for 30 minutes in DMEM supplemented with 0.5% BSA (DMEM/BSA) and incubated for 1 hour with DMEM/BSA containing 20 μg/ml Alexa 647 conjugated transferrin. Cells were incubated for 2 min in DMEM/BSA at 37°C and transferred back on ice. Transferrin at the cell surface was removed by incubating cells for 3 min in ice cold stripping buffer (0.1M Glycine, 150 mM NaCl, pH 3.0), washed twice with ice cold DMEM/BSA and incubated for indicated times at 37°C in DMEM/BSA supplemented with 500 ug/ml unlabelled transferrin. Cells were fixed 10 min with 4% (w/v) paraformaldehyde in PBS. For EEA1 staining, cells were permeabilized for 5 min with 0.1% Triton X-100 (v/v) in PBS and incubated with polyclonal anti-EEA1 antibodies diluted with 3% BSA (w/v) in PBS for 45 min at room temperature. For Rab11 staining, cells were incubated for 45 min with anti Rab11 antibodies diluted with 0.2% saponin (w/v), 1% BSA in PBS. After 3 washes in PBS, primary antibodies were revealed with relevant Alexa Fluor-conjugated secondary antibodies. Cells were mounted in Mowiol 40-88 (Sigma) and observed under TCS SP5 confocal microscope (Leica Microsystems, Nanterre, France) using a 63x objective (NA 1.40).

### Image analysis

In order to quantify Tf uptake, a stack of 3 planes with large field of view (1024×1024 at 700 Hz, 91 nm pixel size, 500 nm z step) were taken by confocal microscopy to handle several cells in one field and to select the best equatorial plane. Images were analyzed using open source Icy software (de Chaumont et al., 2012; http://icy.bioimageanalysis.org/) and protocol (available on Icy website: NewColocalizer_with_binary_and_excel_output_v1_batch). Each cells were manually delineated using phase contrast images and vesicle segmented using wavelet spot detection (Olivo-Marin, 2002). Segmented objects were considered colocalized if the distance between their centroid were below 4 pixels and expressed as a percentage. To estimate the relative amount of Tf in EEA1 or Rab11 compartments, integrated Tf-A647 fluorescence in objects resulting from the segmentation of each compartment was measured and normalized to the total amount of Tf-A647 measured in the cell.

### Flow cytometry analysis

HeLa cells were serum starved for 30 min with DMEM/BSA at 37°C and incubated for 15 min with 10 μg/ml fluorescent transferrin in DMEM/BSA. Cells were transferred on ice, lifted by gentle scraping in PBS containing 0.5 mM EDTA, resuspended and incubated with propidium iodide to exclude dead cells prior to analysis with the Macsquant analyzer (Miltenyi biotec, Bergisch Gladbach, Germany). For kinetics analysis of transferrin recycling, after, cells were incubated for 1 hour at 4°C with 20 ug/ml of fluorescent transferrin following serum starvation. Cells were washed with ice cold Hank’s Balanced Salt Solution (HBSS) and either analyzed by flow cytometry to evaluate transferrin binding to the cell surface or incubated for the indicated amount of time at 37°C before acid wash to remove remaining cell surface transferrin. At least 10000 cells were analyzed and the mean fluorescence intensity of each condition was normalized to the mean fluorescence intensity of the control cells for comparison.

### Western blot

Cells were washed twice with ice cold PBS and lysed in RIPA buffer (20 mM Tris/HCl pH 7.4, 137 mM NaCl, 10% glycerol, 1% Triton X-100, 0.1 % SDS, 0.5 % sodium deoxycholate, 1 mM EDTA) for 20 minutes on ice. Lysates were centrifuged for 10 min at 20 000 g and protein concentration determined using BioRad protein assay. 20 ug of proteins were separated on Novex 4-12% Bis-Tris gel (ThermoFisher Scientific) and transferred to nitrocellulose membrane. Blots were blocked for 1h at room temperature in Tris buffer saline containing 3% BSA and 0.1% Tween-20 and incubated overnight with indicated primary antibodies. After 3 washes, blots were incubated with secondary antibodies coupled to HRP. Detection was carried out with Super Signal West Dura Extended Duration substrate (ThermoFisher Scientific) and immunoreactive bands imaged using Chemismart 5000 (Vilber Lourmat, Marne-la-Vallée, France).

### Time lapse video microscopy

HeLa cells were transfected with GFP-ITSN2-S and TagRFP-T-EEA1 (Addgene, reference: 42635) and observed under spinning disk microscope (Zeiss Axio Observer Z1 with a Yokogawa CSU X1 confocal head) at 37°C using x63 1.4 NA. Each fluorescent protein was sequentially acquired to avoid bleed through using 488 and 561 laser lines at 0.5 Hz. BP525/50 and BP629/62 emission filters were used to retrieve GFP-ITSN2S and TagRFP-EEA1 signals respectively. The acquisition sequence was driven by Metamorph software (Molecular devices, San Jose, CA, USA). Movies were opened in Icy software and single spot detected for each channel and analyzed using Track Manager plugin. The track processor for interaction analysis was used to estimate vesicle association duration. Vesicles were considered associated if they were separated by less than 3 pixels. Only vesicles that remained associated for more than 3 consecutive planes (6 secondes) were considered for analysis.

### Statistical analysis

Statistical analyses were performed using SigmaPlot 11.0 (Ritme, Paris, France). One way ANOVA followed by Holm-Sidak post hoc test was performed to assess the difference between groups when data followed normal distribution and equal variance. If normality test failed, significance was estimated by non parametric Kruskal-Wallis followed by Dunn’s post-hoc test.

---

**Figure S1:**
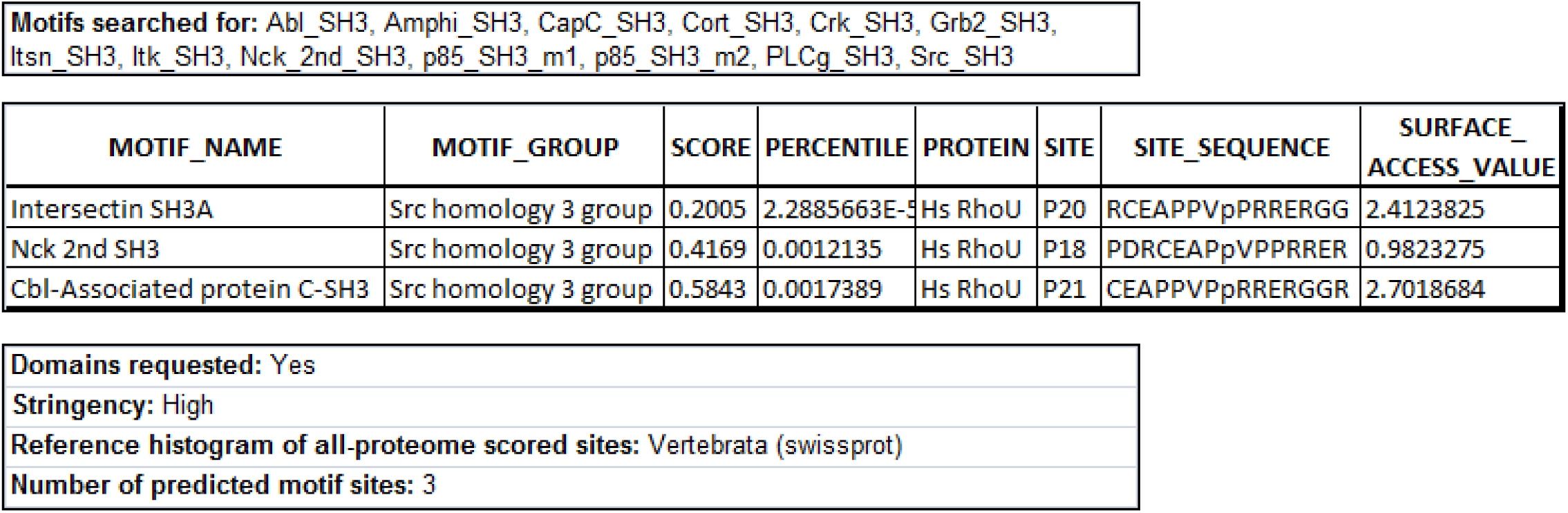
Search for potential SH3 binding sequence in RhoU. The mouse RhoU protein sequence (accession number NP_598716) was searched for potential SH3 binding partners using Scansite database (http://scansite3.mit.edu). The list of motifs searched for is shown and the best hit with high stringency criteria corresponded to the SH3A domain of ITSN1.

